# Single-cell-resolved spatial multi-omics identifies mTOR-driven neuronal and astroglial pathogenicity underlying epileptogenic focal cortical dysplasia

**DOI:** 10.64898/2026.01.23.701175

**Authors:** Kazumi Shimaoka, Satoshi Miyashita, Keiya Iijima, Kaoru Yagita, Nao K N Tabe, Koichi Hashizume, Nariko Arimura, Shuyu He, Minami Mizuno, Kayo Nishitani, Kanako Komatsu, Kumiko Murayama, Masaki Sone, Emi Usukura, Terunori Sano, Shinichiro Taya, Tomoki Nishioka, Kozo Kaibuchi, Tomoo Owa, Masaki Takao, Masaki Iwasaki, Mikio Hoshino

## Abstract

Focal cortical dysplasia type II (FCDII) is a malformation of cortical development caused by somatic mutations in the mTOR signaling pathway. Two hallmark pathological cell types in FCDII, dysmorphic neurons (DNs) and balloon cells (BCs), arise as a result of somatic mutations in the mTOR signaling pathway and are implicated in the pathophysiology of drug-resistant epilepsy. However, how these somatic mutations reshape cell states within the human cortex remains poorly understood.

Here, we integrate imaging-based spatial transcriptomics (iST), single-nucleus RNA sequencing, and proteomics of surgically resected FCDIIb tissue to define the transcriptional and proteomic profiles of DNs and BCs. Spatial mapping of iST data resolved transcriptional signatures in histologically validated DNs and BCs within FCDIIb sections. Integrative omics analysis further revealed that DNs show upregulation of PI3K-AKT-mTOR and p53-CROT metabolic programs accompanied by suppression of synaptic signaling, whereas BCs exhibit transcriptional signatures of reactive astrocytes with increased phagocytic and immune-like activity.

These data delineate cell-type-specific consequences of somatic mTOR pathway mutations at single-cell resolution and reveal previously unrecognized metabolic and immunoregulatory mechanisms contributing to epileptogenesis in drug-resistant epilepsy.

Our study establishes a spatial multi-omics framework for dissecting human cortical malformations and highlights potential therapeutic targets for drug-resistant epilepsy.

## Main

Focal cortical dysplasia type II (FCDII) is a malformation of the cerebral cortex and is a leading cause of drug-resistant epilepsy in children^1^. As antiepileptic drugs are largely ineffective for FCDII, surgical resection of the epileptogenic lesion remains an effective treatment to reduce seizures^2^. Histopathologically, FCDII is characterized by disrupted cortical lamination, a blurred gray-white matter boundary, and the presence of pathological cell types, including dysmorphic neurons (DNs) and balloon cells (BCs)^3^. DNs, derived from excitatory neurons, are hallmark histological features of both FCDIIa and IIb, whereas BCs, of astrocytic origin, are uniquely associated with FCDIIb.

DNs exhibit aberrant activation of the mTOR signaling pathway due to somatic mutations in mTOR-related genes and reduced action-potential firing with diminished glutamatergic inputs^4–6^. Intriguingly, DN-rich areas are hyperexcitable^7,8^, implicating the function of DNs in epileptogenesis; however, the molecular mechanisms underlying physiological properties of DNs remain poorly understood. In contrast, BCs are generally considered electrically silent, express stem cell-associated markers, and lack detectable electrophysiological activity^9^, yet their precise role in epileptogenesis also remains unclear. Thus, although DNs and BCs are central to the pathology of FCDIIb, their molecular characteristics and pathogenic functions are incompletely understood.

Here, we combined imaging-based spatial transcriptomics (iST) with single-nucleus RNA sequencing (snRNA-seq) to define the transcriptional profiles of DNs and BCs identified by morphology and marker expression in human FCDIIb tissue. Integrative analysis revealed molecular profiles and hallmark signaling pathways in DNs and BCs. Complementary proteomic profiling of FCDIIb brain tissue identified candidate marker proteins for each cell type, providing further mechanistic insight into epileptogenesis in FCDIIb.

## Results

### Study design overview

To approach the molecular characteristics of FCDIIb brain at a single-cell resolution, we analyzed five FCDIIb patients using multi-modal omics-based profiling, including imaging-based spatial transcriptomics (iST) analysis, snRNA-seq, and proteomics (Fig. 1a, Table S1). Neurotypical cortical regions from hippocampal sclerosis (HS) and glioneuronal hamartoma patients were used as controls. Genome analysis of FCDIIb samples identified three types of somatic mutations in *MTOR* gene from three patients and one germline mutation in *TSC1* gene from one patient (Table S1). All detected mutations in *MTOR* were previously reported pathogenic mutations known to increase the mTOR signaling pathway activity^4,10–12^.

**Figure 1.**
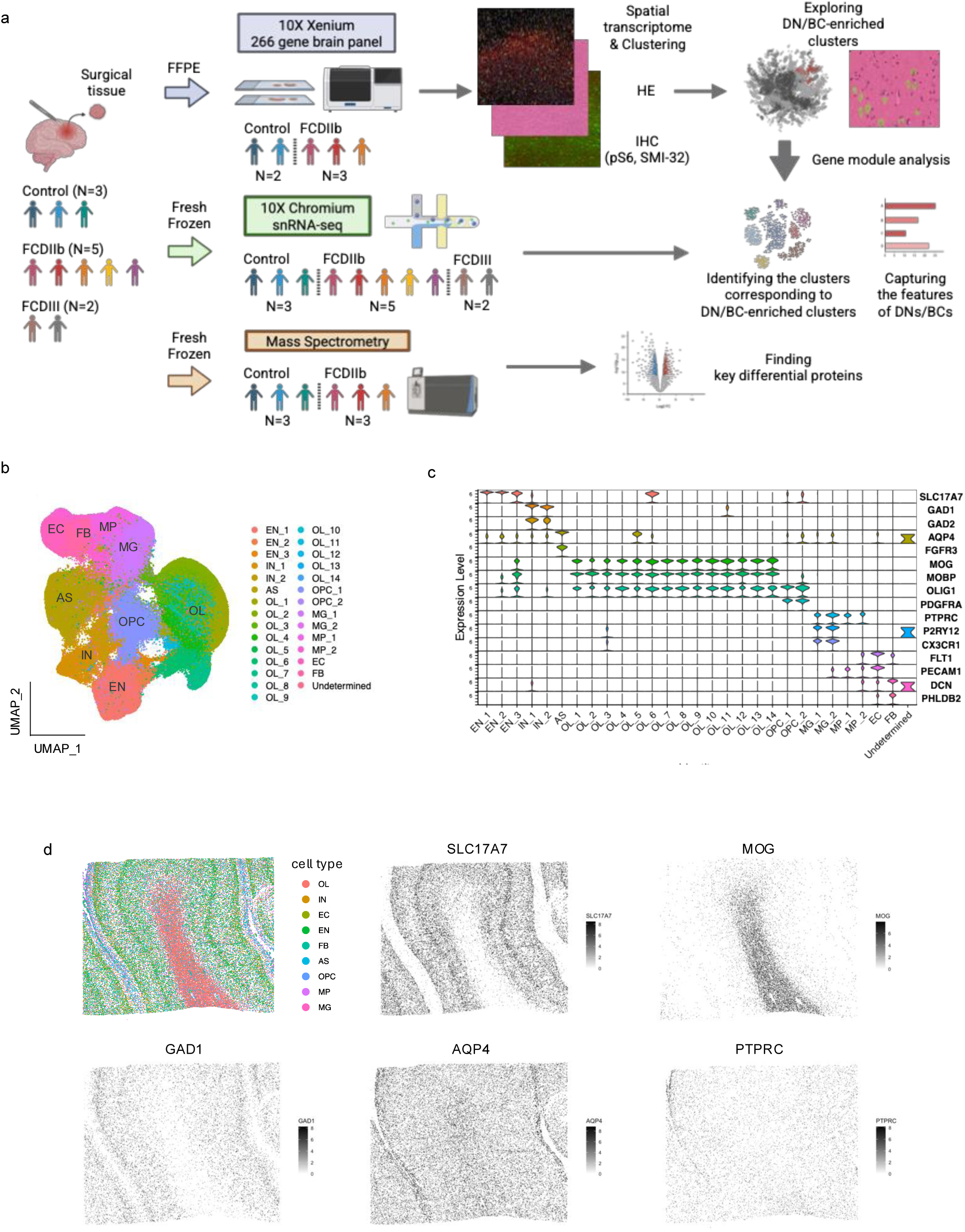
Overview of Xenium spatial transcriptomic workflow and cell-type annotation. (a) Experimental design and analytical workflow. (b) Uniform manifold approximation and projection (UMAP) embedding of iST data showing major cell classes, including excitatory neurons (EN), inhibitory neurons (IN), astrocytes (AS), oligodendrocytes (OL), oligodendrocyte precursor cells (OPC), microglia (MG), macrophages (MP), endothelial cells (EC), and fibroblast (FB). (c) Violin plots showing expression of canonical marker genes across clusters, confirming cell type identities (e.g., *SLC17A7* for EN, *GAD1/GAD2* for IN, *AQP4/FGFR3* for AS, *MOG/MOBP* for OL, *PDGFRA* for OPC, *PTPRC* for immune cells, *P2RY12/CX3CR1* for MG, *PECAM1/FLT1* for EC, *DCN* for FB). (d) Spatial mapping of annotated cell types and feature plots showing spatial gene expression patterns of representative markers (EN:*SLC17A7*, OL:*MOG*, IN:*GAD1*, AS:*AQP4*, MG:*PTPRC*).

### Single-cell resolution spatial transcriptomics of FCDIIb brain

To investigate the transcriptomic features of FCDIIb, we first performed iST analysis using the Xenium platform, profiling 266 genes across two controls, three FCDIIb brain sections harboring *MTOR* mutations and one peripheral section located next to the lesion site of FCDIIb (patient 2, Table S2). We confirmed that DNs and BCs were observed in all FCDIIb sections used for iST analysis but not in the control sections and the peripheral section. After iST data acquisition, we performed immunostaining with anti-phospho-S6 (pS6) and SMI-32 antibodies, as well as hematoxylin-and-eosin staining on the same slides following iST analysis (Fig. 1a), enabling direct comparison of transcriptomic profiles with marker protein expression and cellular morphology.

Gene expression was quantified per cell based on segmentation of nuclear (DAPI) and cell soma staining (18S ribosomal RNA and VIMENTIN (VIM), see also Method). After quality control filtering, 880,831 cells (247,095 from control, 470,417 from FCD, 163,319 from peripheral) were retained and classified into 29 clusters, composed of excitatory neuron (EN), inhibitory neuron (IN), astrocyte (AS), oligodendrocyte (OL), oligodendrocyte precursor cell (OPC), microglia (MG), macrophage (MP), endothelial cell (EC), and fibroblast (FB), according to the expression of cell-type-specific markers (Fig. 1b,c). All cell types were detected across all sections from FCDIIb, peripheral region, and control samples, although excitatory neurons were moderately reduced in patient 2 (Fig. S1a). Spatial mapping of annotated cell types in control and peripheral region sections clearly visualized the boundary structure between the gray matter and white matter, which was further supported by the spatial distribution of cell type-specific marker gene expression (*SLC17A7*: Excitatory neuron EN, *MOG*: Oligodendrocyte OL, *GAD1*: Inhibitory neuron IN, *AQP4*: Astrocyte AS, *PTPRC*: Microglia MG) (Fig.1d, Fig.S1b). By contrast, the boundary structures were blurred in FCDIIb sections, consistent with the pathohistological feature of FCDII (Fig. S1b).

### Identification and characterization of DNs and BCs in iST analysis

DNs and BCs are hallmark histopathological features of FCDIIb, characterized by enlarged cell bodies and activated mTOR signaling. A recent transcriptomic analysis of microdissected DNs and BCs suggested that DNs and BCs have excitatory neuron- and astrocyte-like identities, respectively^13^. We first validated the cell types of DNs and BCs by mapping cluster annotations (Fig. 1b) onto the histological space (Fig. S2a, b). We found that phosphorylated-S6-, a marker for mTOR signaling activation, and SMI32-, marker for DNs, positive DNs mapped to the EN cluster, whereas pS6- and VIM/αSMA-positive BCs mapped to the AS cluster(Fig. S2a, b). Hematoxylin and eosin (H&E) staining further confirmed the morphological features of DNs and BCs (Fig. S2a, b).

### Determination of DN-enriched cluster by single-cell resolution spatial transcriptomics

To investigate the molecular profiles of DNs, we performed sub-clustering of excitatory neurons in EN cluster (Fig. 2a) and identified 17 sub-clusters based on marker expression and spatial distribution (Fig. 2a-c). In control sections, excitatory neuron subclusters (L2, L2-3, L4, L4-5, L5_1, L5_2, L5_3, L5-6_1, L5-6_2) displayed a layered structure (Fig. 2c), reflecting distinct transcriptomic signatures corresponding to cortical layer identities. In FCDIIb sections, the sub-clusters exhibiting layer-specific gene expression were maintained, but the distribution of neurons within each sub-cluster was remarkably broader than control, partially disrupting the layer structure (Fig. 2d). In addition, sub-clustering analysis revealed FCDIIb-specific sub-clusters (FCDIIb lesion-enrich_1,2,3), whose neurons were widely distributed across the gray matter (Fig. 2d). We further found that neurons in lesion enrich_1 and enrich_2 expressed both upper- and deep-layer neuronal markers, whereas those in lesion enrich_3 expressed only deep-layer markers (Fig. 2b). Lesion enrich_1 was predominant in patient 1 and 3, lesion enrich_2 in patient 1 and 2, and lesion enrich_3 cells were detected in small numbers in patient 3 (Fig. S2c). In the peripheral section of FCDIIb, although layer structures were maintained, specific clusters showing upper-(non lesion enrich_1) or deep-layer (non lesion enrich_2) transcriptomic features were observed (Fig. 2b,e). Consistent with their transcriptomic features, enrich_1 cells were located around L2-3 region, whereas enrich_2 cells around L5-6 region (Fig. 2e). FCDIIb lesion enrich_1, 2, and 3 cells, observed exclusively in FCDIIb sections, were absent in control and peripheral sections (Fig. S2d,e), suggesting that these excitatory neurons were candidates for DNs. H&E staining of FCDIIb enrich_1, 2, and 3 cells confirmed the enlarged cell bodies (Fig. 3a), and immunostaining showed strong pS6 and SMI32 signals were observed in FCDIIb lesion enrich_1, 2, and 3 cells (Fig. 3b-d), suggesting that DNs were enriched in these clusters.

**Figure 2.**
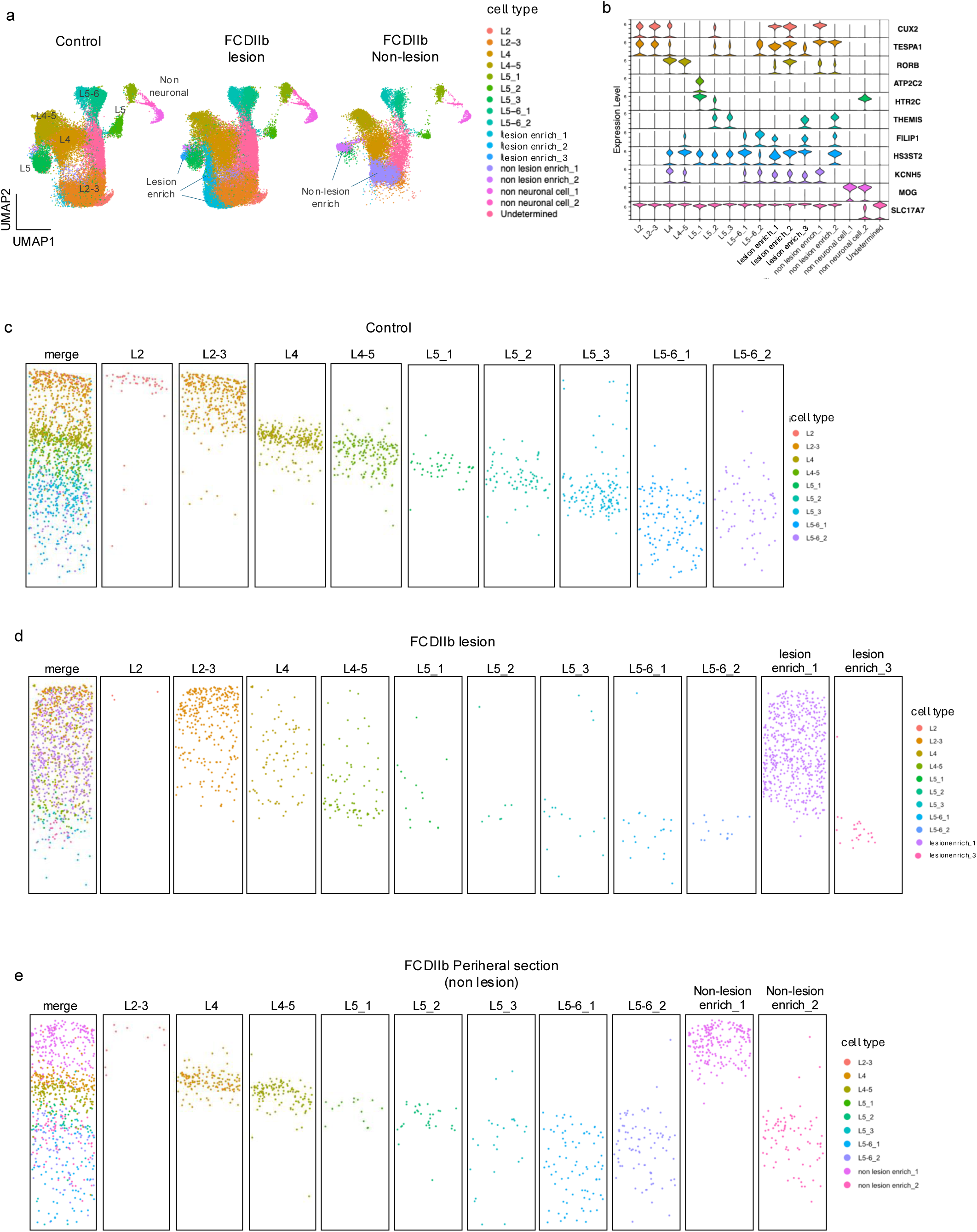
Spatial mapping of layer specific subclusters and lesion-enriched neuronal populations. (a) UMAP embedding of neuronal clusters from control, FCDIIb lesion, and FCDIIb non-lesional peripheral regions, highlighting layer-specific excitatory neuron populations and lesion-enriched neuronal clusters. (b) Violin plot showing expression of representative marker genes indicating layer identity and lesion-enriched populations. *MOG* expression revealed non-neuronal cells that were misclassified in the original clustering and were subsequently separated during subclustering. (c-e) Spatial mapping of annotated cells showing layer-specific excitatory neuron subclusters across control tissue (c), FCDIIb lesion (d), peripheral non-lesion (e) sections.

**Figure 3.**
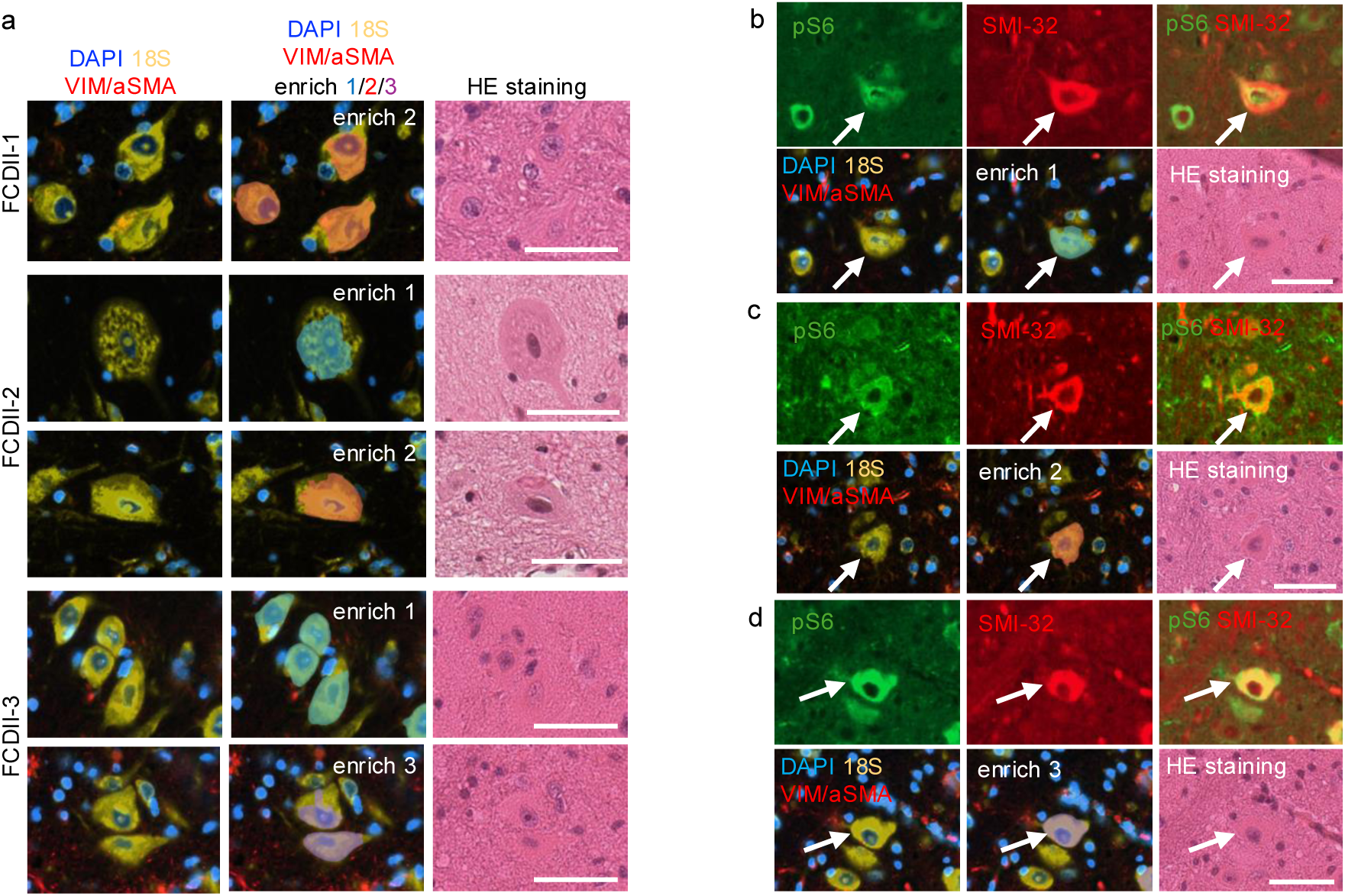
Validation of DNs by immunofluorescence and histology. (a) Representative immunofluorescence images from FCDIIb cases showing DN-enriched cellular regions labeled with DAPI and 18S RNA, and reactive/structural markers (VIM/αSMA), alongside corresponding H&E staining. Panels include representative fields from multiple patients and multiple enrichment regions. (b-d) pS6 and SMI32 expressions in cells from FCDIIb lesion-enriched clusters (enrich 1-3). pS6 and SMI32 signals were observed in cells from FCDIIb enrich_1 from patient 3 (b), FCDIIb enrich_2 from patient 1 (c), and FCDIIb enrich_3 from patient 3 (d) cells. Scale bars: 50 µm

### Determination of BC-enriched cluster by single-cell resolution spatial transcriptomics

To investigate the molecular characteristics of BCs, we performed sub-clustering of astrocytes from control, FCDIIb, and peripheral region sections. Sub-clustering divided astrocytes into 16 subclusters and identified two FCDIIb-enriched clusters (FCDIIb enrich_1 and 2) which were detected from FCDIIb lesional and peripheral sections and not from control sections (Fig. 4a-c). Cells in FCDIIb enrich_1 were predominantly enriched in the FCDIIb lesional sections, whereas enrich_2 cells were observed in both lesional and peripheral sections (Fig.4c, Fig.S3a,b). Thereby, we compared the cell morphologies of FCDIIb enrich_1 and FCDIIb enrich_2 cells, revealing that FCDIIb enrich_1 cells manifested the hallmark morphological features of BCs with enlarged cell bodies, while FCDIIb enrich_2 cells in lesional and peripheral sections resembled the morphology of reactive astrocytes showing short and rugged processes (Fig. 4d,e). Moreover, strong pS6 signal was observed in FCDIIb enrich_1 cells, indicating the activation of mTOR signaling in FCDIIb enrich_1 cells. These findings suggested that BCs were enriched in the FCDIIb enrich_1 of astrocyte subclusters (Fig. 4f). FCDIIb enrich_2 cells also showed strong signal of pS6 (Fig.4g), indicating that mTOR signaling was also activated in FCDIIb enrich_2 cells.

**Figure 4.**
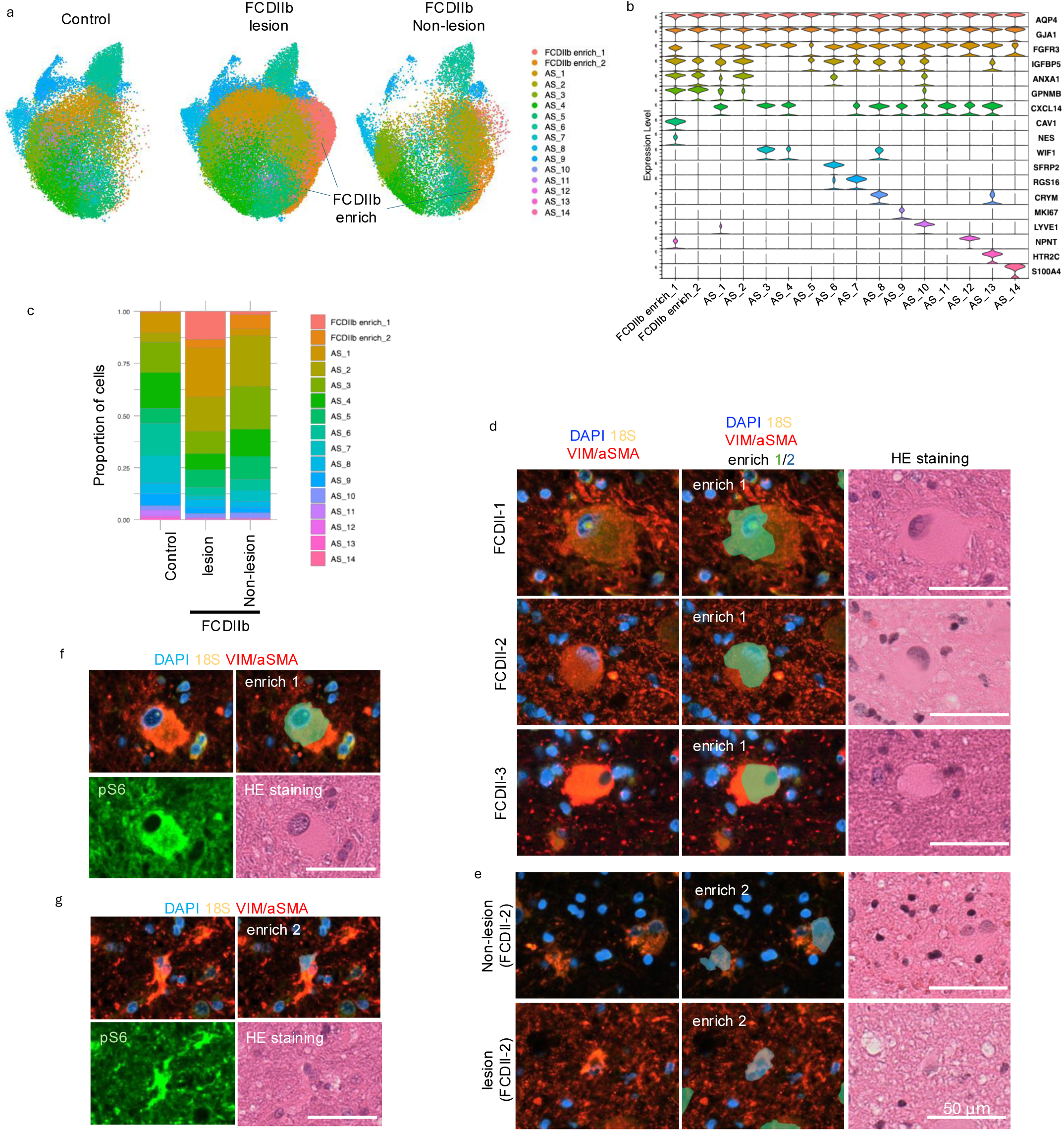
Identification of BCs by iST analysis in FCDIIb sections. (a) UMAP embedding of astrocyte clusters across control, FCDIIb lesion, and FCDIIb non-lesion regions. Lesion-enriched astrocyte subclusters were highlighted. (b) Violin plot showing astrocyte markers and lesion-associated genes (*AQP4, GJA1, FGFR3, IGFBP5, ANXA1, GPNMB, CXCL14, CAV1, NES, WIF1, SFRP2, RGS16, CRYM, MKI67, LYVE1, NPNT, HTR2C, S100A4*). (c) Proportional composition of astrocyte subclusters across groups/regions. (d,e) Morphology and marker expression of lesion-enriched astrocyte subclusters. H&E staining and boundary staining (DAPI, 18S, and VIM/αSMA) in FCDIIb enrich_1 (d) and 2 (e). FCDIIb enrich_1 cells observed in patient 1, 2 and 3 exhibited large cell body, whereas FCDIIb enrich_2 cells had small cell size. (f,g) pS6 immunosignal, cell boundary staining (DAPI, 18S, and VIM/αSMA) and H&E in FCDIIb enrich_1 cell from patient 1 (f) and FCDIIb enrich_2 from lesion site of patient 2 cell (g). Scale bars: 50 µm

Taken together, combining sub-clustering and immunostaining of marker proteins of DNs or BCs successfully identified histologically defined DNs and BCs in FCDIIb.

### Identification of prospective DN and BC clusters in snRNAseq of FCDIIb

To overcome the limitation of the numbers of genes detectable by Xenium iST analysis and to characterize the molecular profiles of DNs and BCs, we next performed snRNA-seq analysis and projected iST-derived DN and BC signatures onto the snRNA-seq space. Five FCDIIb, and three controls from neurotypical regions of HS and glioneuronal hamartoma were enrolled for snRNA-seq analysis. Two temporal lobe neocortical regions of HS patients with cortical dyslamination (FCDIIIa) were also included in the analysis as negative controls lacking DNs and BCs. After quality control, 22,491 nuclei were finally retained and annotated as excitatory neuron (EN), inhibitory neuron (IN), astrocyte (AS), oligodendrocyte (OL), oligodendrocyte precursor cell (OPC), microglia (MG), macrophage (MP), endothelial cell (EC), or fibroblast (FB, Fig.5a,b). We confirmed that all cell type clusters were detected across samples (Fig.5c). We first performed sub-clustering of excitatory neurons and identified layer specific pyramidal cortical neuron clusters defined by canonical layer marker genes, and two FCDIIb enriched clusters (Fig 5d, S4a). To confirm whether FCDIIb enriched clusters in snRNA-seq space show similar molecular characteristics of DNs revealed by iST analysis, we extracted the 13 differentially expressed genes of FCDIIb lesion enrich_1 and _2 clusters in iST space (Fig. 2a, 3a, DN module) and calculated the module score of DN module (Fig. 5e, see also Methods). Among excitatory neuron clusters, one cluster exhibited significantly high score of DN modules (the prospective DN cluster), suggesting that DNs were enriched in this cluster (Fig. 5f, g). The prospective DN cluster was predominantly detected in FCDIIb samples (Fig. 5h). We next examined the expressions of known DN markers^13^. Expressions of *NEFM, NEFH, STMN2, SLC1A1, and TUBB3* showed significant increase in prospective DN cluster compared to the other excitatory neuron clusters (Fig. 5i). *NEFL and NME7* genes showed moderately high expressions, whereas *PROM1 and MT1G* were barely detected in both prospective DN and the other clusters of our dataset. These results indicate that the prospective DN cluster identified in snRNA-seq was highly enriched for nuclei of DNs.

**Figure 5.**
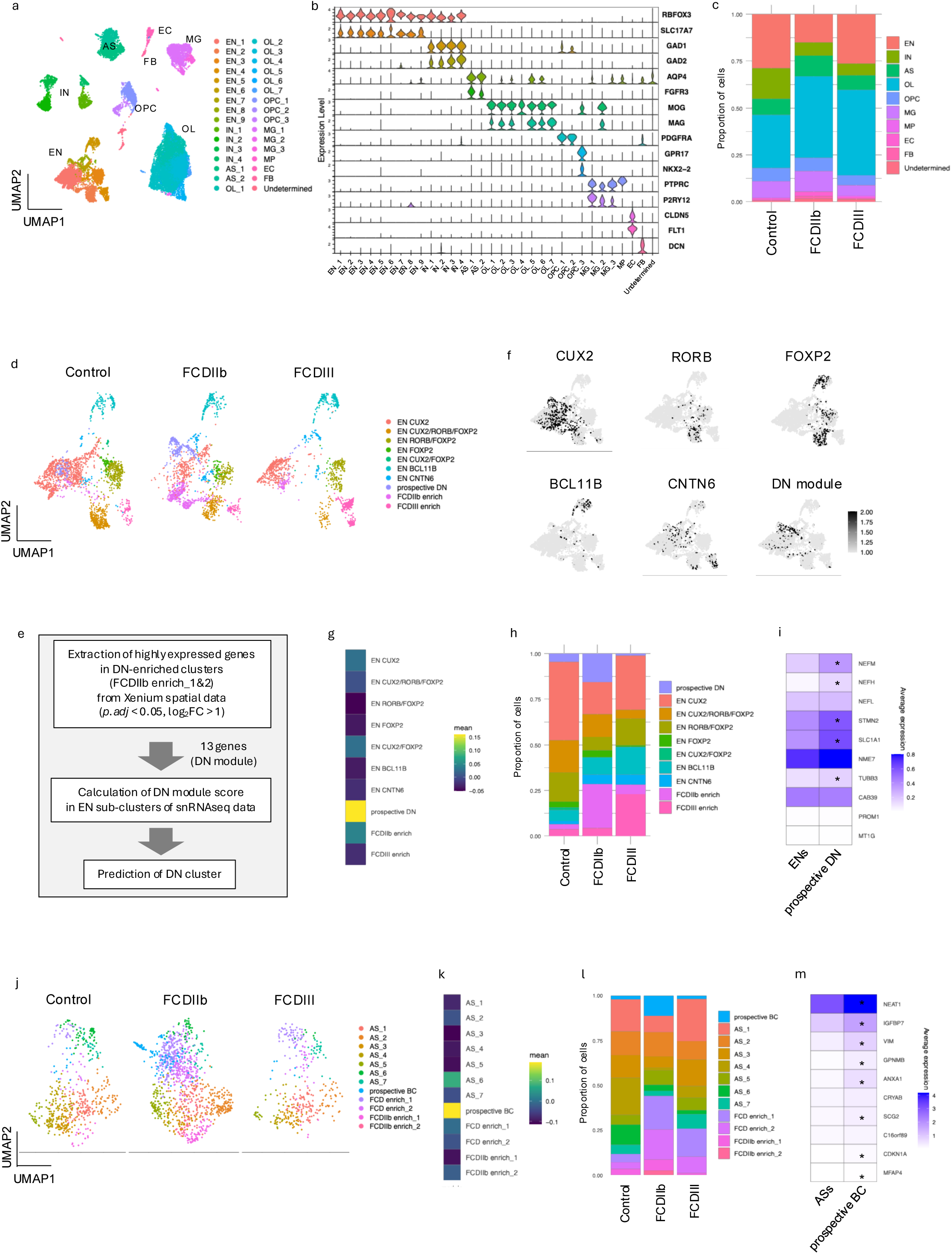
Projection of iST-derived modules onto snRNA-seq space to identify DN and BC clusters. (a) UMAP embedding of snRNA-seq data showing major cell classes, including excitatory neurons (EN1-9), inhibitory neurons (IN1-4), astrocytes (AS-1,2), oligodendrocytes (OL1-7), oligodendrocyte precursor cells (OPC1-3), microglia (MG1-3), macrophages (MP), endothelial cells (EC), fibroblast (FB), and Undetermined. (b) Violin plots of major cell type marker expressions confirming cluster identity across clusters. (c) Proportional composition of cell-type across control, FCDIIb, and FCDIII samples. (d) UMAP embedding of excitatory neuron (EN) subclusters separated by condition (Control, FCDIIb, FCDIII). (e) Analytic workflow for DN module gene extraction from DN-enriched spatial clusters (FCDIIb enrich_1/2) and module scoring in EN subclusters using snRNA-seq data to predict DN-associated neuronal clusters. (f) Feature plots showing expression of layer specific EN markers (CUX2, RORB, FOXP2, BCL11B, CNTN6) and DN module score. (g) Visualization of DN module score across EN subclusters. (h) Proportional composition of EN subclusters across control, FCDIIb, and FCDIII samples. (i) Expression of known DN markers in prospective DN compared with the other excitatory neurons. *Adjusted p value < 0.05. (j) UMAP embedding of astrocyte (AS) subclusters separated by condition (Control, FCDIIb, FCDIII). (k) Visualization of BC module score across AS subclusters. (l) Proportional composition of EN subclusters across control, FCDIIb, and FCDIII samples. (m) Expression of known BC markers in prospective DN compared with the other astrocytes. *Adjusted p value < 0.05.

Next, we performed sub-clustering of astrocyte cluster from snRNA-seq data (Fig. 5j, S4b), and assessed a BC module comprising 22 iST-derived genes from the FCDIIb enrich_1 cluster (Fig. 4a). We identified one cluster exhibiting significantly high expression of BC module score and predominantly observed in FCDIIb (Fig. 5k,l, S4c). Thus, we named this cluster as prospective BC cluster. We also found that specific subclusters (FCDIIb enrich_1/2) exclusively observed in FCDIIb samples and subclusters (FCD enrich_1/2) observed both FCDIIb and FCDIIIa samples (Fig. 5j, l). We also confirmed the eight out of ten known BC markers^13^ *(NEAT1*, *IGFBP7*, *VIM*, *GPNMB*, *ANXA1*, *SCG2*, *CDKN1A*, and *MFAP4*) showed significantly high expressions in the prospective BC cluster compared to the other clusters of astrocytes, suggesting that nuclei of BCs were enriched in this cluster (Fig. 5m). Furthermore, the prospective BC cluster showed high expression of reactive astrocyte markers (*C3*^14^, *CD44*^14^*, MAOB*^15^), suggesting that BCs shared transcriptional features with reactive astrocytes (Fig. S4b,c). Together, projecting iST-derived gene module expression onto snRNA-seq space enabled the robust identification of DNs and BCs, expanding the number of detectable genes from 266 in iST to over 10,000 in snRNA-seq.

### Characterizing molecular features of prospective DNs and BCs

We further characterized the molecular features of DNs and BCs identified in snRNA-seq space. Differential expression analysis between prospective DNs and the other excitatory neuron clusters detected 1,434 significantly upregulated differentially expressed genes (DEGs) and 104 significantly downregulated DEGs (p.adj < 0.05, |log_2_FC| > 1; Table S3). KEGG and reactome analysis using upregulated DEGs identified “PI3K-Akt signaling pathway” and “p53-p21 signaling” as a significantly altered pathway, consistent with activation of mTOR signaling pathway in DNs and the senescence feature of DNs^7,16^ (Fig. 6a, Fig. S5a, Table S4,5). These results confirmed that the prospective DN cluster recaptured the known features of DNs. In addition, KEGG analysis identified “Focal adhesion” and “Cell adhesion molecules” related genes in the upregulated DEGs (Fig 6a, Table S4). Gene ontology (GO) analysis of biological processes demonstrated that “extracellular matrix (ECM) organization” and “MAPK cascade” related genes were enriched among the upregulated DEGs (Fig 6b, Table S6). In contrast, synapse-related GO terms (“synaptic plasticity”, “positive regulation of synaptic transmission”, and “long-term synaptic potentiation”) and synapse-related KEGG pathways (“Neuroactive ligand signaling”, “GABAergic synapse”, and “Synaptic vesicle cycle”) were enriched in the downregulated DEGs, suggesting the impairment of synaptic activity in DNs exhibiting abnormal electrophysiological characteristics^5^ (Fig. 6c and S5b, Table S7).

**Figure 6.**
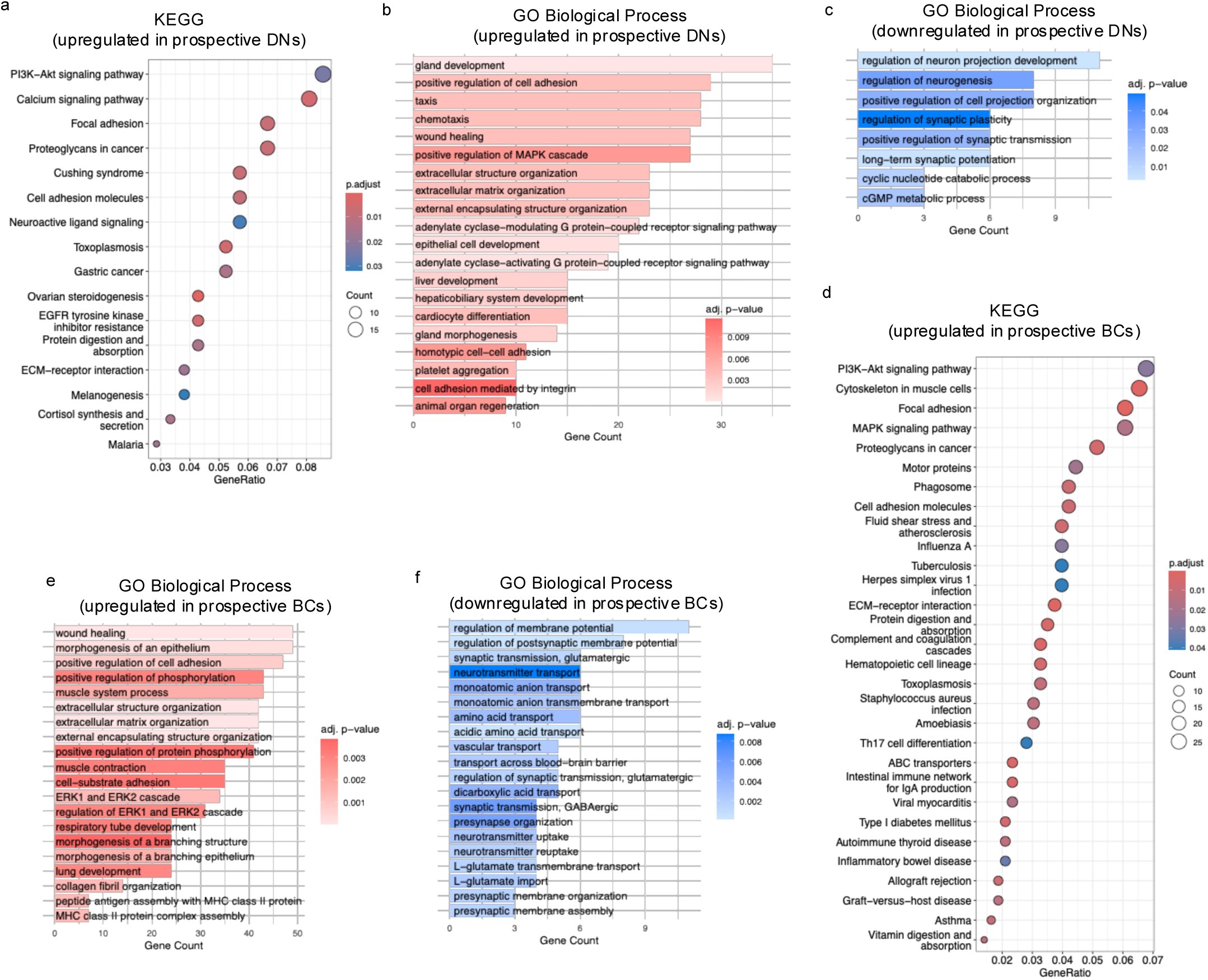
Functional enrichment analyses of prospective DNs and BCs. (a) KEGG pathway enrichment for genes upregulated in prospective DN cluster. (b) GO Biological Process enrichment of genes upregulated in prospective DN cluster. (c) GO Biological Process enrichment of genes downregulated in prospective DN cluster. (d) KEGG enrichment of genes upregulated in prospective BC cluster. (e) GO Biological Process enrichment of genes upregulated in prospective BC cluster. (f) GO Biological Process enrichment of genes downregulated in prospective BC cluster.

Next, we performed differential expression analysis between prospective BC cluster and the other astrocyte clusters, identifying 2,078 significantly upregulated DEGs and 98 significantly downregulated DEGs (p.adj < 0.05, |log_2_FC| > 1; Table S8). KEGG analysis for the upregulated DEGs of prospective BC cluster also identified “PI3K-Akt signaling pathway” as an enriched pathway (Fig. 6d, Table S9), consistent with the previously reported activation of mTOR signaling pathway in BCs^17^. Notably, KEGG analysis identified phagosome-related pathway with significantly upregulated DEGs (Fig. 6d, Table S10). In addition, MAPK signaling pathway, and ABC transporter pathways, both associated with phagocytosis^18,19^, were identified as upregulated pathways (Fig. 6d, Table S10). Consistently, GO analysis also detected the increase of ERK1/2 signaling (Fig.6e, Table S11) which is known to regulate phagocytosis activity^20,21^. Recently it has been suggested that astrocytic phagocytosis was important to maintain the brain homeostasis in the normal and disease brain^22,23^, and phagocytosis activity was promoted by PI3K and mTOR signaling activity^24,25^. Thus, our results suggested that BCs might exhibit enhanced phagocytosis activity. Moreover, MHC class II protein assembly-related pathways were also identified in upregulated DEGs (Fig.6e, Table S11), suggesting that BCs might exhibit properties of immune cell-like astrocytes as proposed previously^26^. Conversely, “synaptic transmission” and “neurotransmitter transport” related GO terms were identified from downregulated DEGs (Fig. 6f, Table S11), suggesting the impairment of normal astrocytic function, including neurotransmitter transport and modulation of synaptic activity^27,28^.

### Identification of differentially expressed proteins in FCDIIb specimens by proteomic analysis

To further explore the molecular features of DNs and BCs at the protein-level, we performed liquid chromatography with tandem mass spectrometry (LC-MS/MS) on surgically resected fresh-frozen brain specimens from three FCDIIb patients and three controls (Fig. 1a). All samples were derived from the same individual with snRNA-seq and iST analysis. In total, 5,737 proteins were detected, and differential expression analysis identified 220 upregulated differentially expressed proteins (DEPs) and 296 downregulated DEPs (Fig. 7a, Table S12). NEFM, NEFH and VIM, previously reported as increased genes in DNs and BCs^29^, were detected among the upregulated DEPs, indicating that proteins derived from DNs and BCs were enriched in our proteomics data.

**Figure 7.**
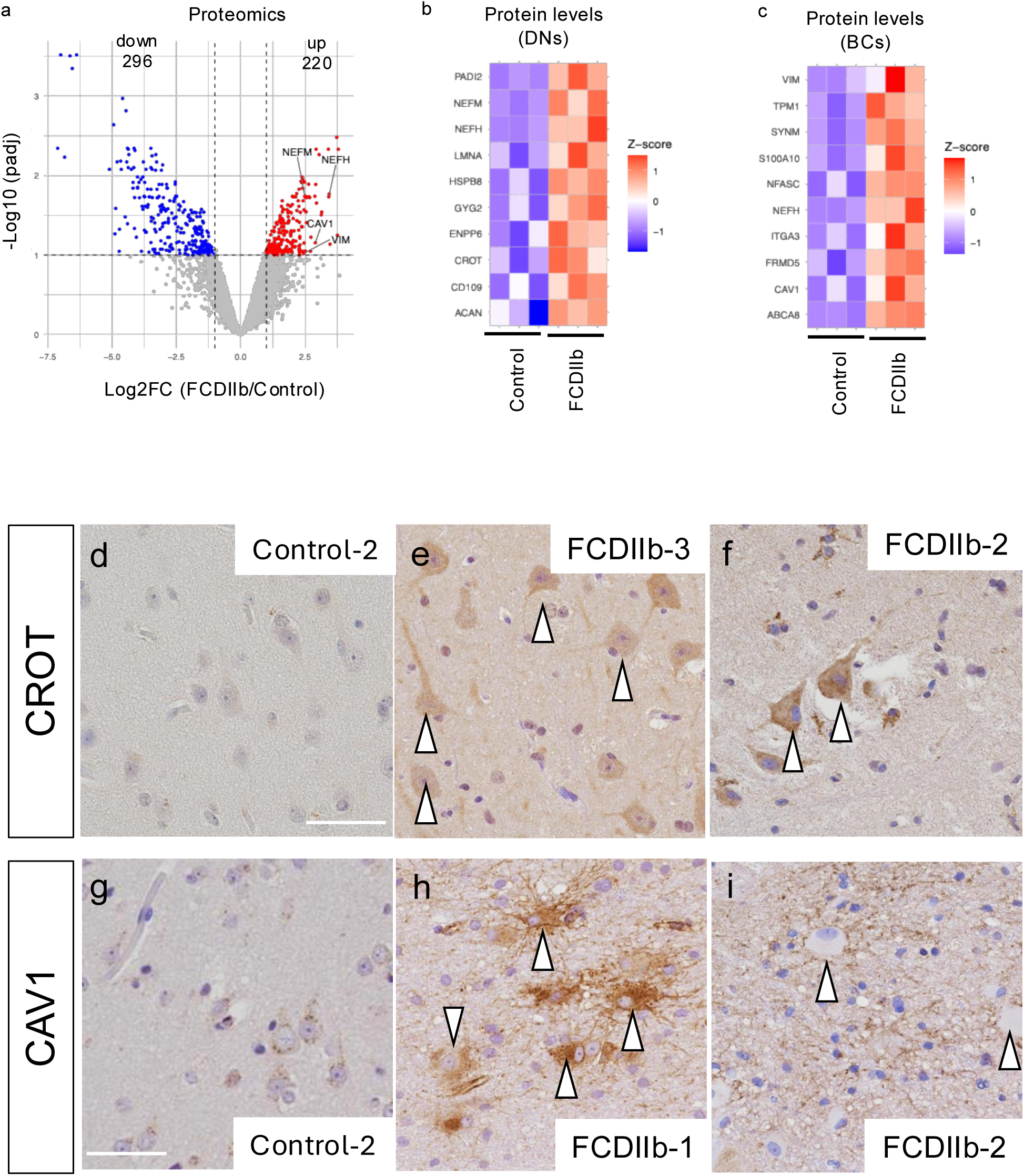
Proteomic validation of lesion-associated DN and BC molecular features. (a) Volcano plot showing differentially expressed proteins between FCDIIb and control samples (log2 fold change vs. -log10 adjusted p-value). (b) Heatmap showing relative protein expression changes in candidate DN-associated proteins across control and FCDIIb samples. (c) Heatmap showing relative expression of candidate BC-associated proteins across control and FCDIIb samples. (d-f) Immunohistochemistry validation of CROT expression across neurons in control tissue and DNs in FCDIIb tissues. Arrowheads indicate DNs. (g-i) Immunohistochemistry validation of CAV1 expression across control tissue and BCs in FCDIIb tissues. Arrowheads indicate BCs.

To link proteomics findings with transcriptomic data, we compared DEPs with DEGs from the prospective DN or the prospective BC clusters identified in snRNA-seq analysis. Finally, 19 proteins were identified as candidate upregulated proteins in DNs and BCs (Fig. 7b,c). Among the 19 genes, we found that *CROT* was mostly expressed in the DNs of snRNA-seq space (Fig. S6a). *CROT* encodes Carnitine O-Octanoyl transferase regulating the fatty acid metabolism^30^, and is a transcriptional target of p53, a well-known senescence marker highly expressed in DNs^7^. In the neurotypical cortical region, CROT expression was restricted to the astrocytes localized beneath the meninges, whereas CROT expression was observed in the DNs as well as the astrocytes in the cortex of FCDIIb (Fig. 7d-f). CROT was rarely detected in other types of neurons (Fig. S6a), indicating that CROT is a novel marker for DNs.

Among the upregulated proteins of BCs, we focused on the Caveolin 1 (CAV1), a principle component of caveolae structure to control the endocytosis^31,32^, as *CAV1* was exclusively expressed in the BC cluster of iST analysis and prospective BC cluster of snRNA-seq among AS clusters (Fig. 4b, Fig. S6b). Although strong CAV1 expression was limited to the vascular cells in neurotypical regions of control sample, CAV1 expression was also observed in BCs of FCDIIb sections (Fig. 7g-i). Notably, CAV1 was localized to the complex processes of BCs. Since recent studies highlighted the involvement of CAV1 in phagocytosis^33,34^, these results further supported the possible function of BC in phagocytosis.

## Discussion

FCDII is a leading cause of drug-resistant epilepsy in pediatric patients, characterized histopathologically by cortical layer disruption and the presence of pathological cells, including DNs and BCs^1,3^. Although recent genomic studies have revealed that somatic mutation in PI3K-AKT-mTOR pathway underlies the pathology of FCDII^10–12^, the molecular and cellular mechanisms driving epileptogenesis remain elusive.

DNs are located within cortical regions where neuronal hyperexcitabilities are observed, yet individual DNs themselves exhibit lower electrophysiological activity than neighboring pyramidal neurons^5,6^. Consistent with this, reduced neuronal activities were also reported in mouse models of FCDII^35^. Moreover, Xu et al. demonstrated that ablating DNs suppresses spontaneous seizures in FCDII mouse model^36^. Together, these findings suggested the contribution of DNs to the epileptogenesis and highlighted them as a potential therapeutic target in drug resistant epilepsy. Defining the molecular features of DNs is, therefore, essential for elucidating cellular mechanisms of epileptogenesis and for developing therapies for FCDII. Recent multimodal-omics studies have begun to characterize DNs: integration of snRNA-seq with single-cell ATAC seq identified several novel DN markers^29^, and simultaneous single-cell genotyping with transcriptome profiling delineated the transcriptomic features of pathogenic cells harboring MTOR mutations^13^. Sequence-based spatial transcriptomics also provided spot level transcriptomic profiles^37,38^. However, capturing molecular profiles of DNs at single-cell resolution has remained challenging. Here, we utilized iST analysis using the Xenium platform, which enables subcellular level detection of transcripts. Our analysis identified transcriptomic features of histologically validated DNs, and integration with snRNA seq expanded the detectable gene set from 266 targeted transcripts to more than 20,000 genes.

We confirmed activation of the PI3K-AKT-mTOR pathway in DNs and observed upregulation of p53 signaling, consistent with previously reported senescence-like features of DNs^7^. Moreover, previously reported DN marker genes were significantly enriched, validating that the putative DN cluster in our analysis is predominantly composed of DN nuclei. We also uncovered downregulation of synaptic plasticity and neurotransmission-related genes in DNs, suggesting a molecular basis for the low firing properties of DNs compared to other neurons^5,6^. In addition, we identified the elevated expression of CROT in DNs by transcriptomic, proteomic and immunostaining analysis. CROT, a peroxisomal enzyme regulating very long-chain fatty acids (VLCFA) metabolism and a transcriptional target of p53, has been shown to promote the cell survival under nutrient-starved condition by increasing the mitochondrial respiration^39^. Recent studies further suggested that VLCFA synthesis influences the neural polarity through the formation of lipid rafts^40^. Thus, aberrant CROT protein expression in DNs may alter the cellular metabolic status contributing to the abnormal morphology, neural activity, and potential cellular resilience to excitotoxic stress after seizures. Collectively, these findings have refined the view that DNs are transcriptionally and metabolically abnormal neurons exhibiting abnormal neuronal activity, destabilizing the cortical network and contributing to the epileptogenesis.

In contrast, BCs have been considered functionally ambiguous, despite their prominence as a histopathological feature in FCDIIb and other mTORopathies such as tuberous sclerosis complex and hemimegalencephaly^41,42^. In this study, we confirmed that BCs displayed the molecular features of reactive astrocytes, including the marker of reactive astrocytes (*MAOB*, *CD44*) and enrichment of immune-related genes. The emerging role of reactive astrocytes in neurological disorders, including Alzheimer’s disease and Huntington’s disease, has been increasingly recognized^43^. Recent studies suggested that reactive astrocyte plays critical roles in epileptogenesis^44,45^, raising the possible involvement of BCs in this process. Interestingly, we observed significant enrichment of phagosome-related pathways in BCs as a significantly upregulated pathway along with abundant expression of CAV1, a regulator of phagocytosis^33,46^, suggesting that BCs may possess increased phagocytotic capacity similar to phagocytic astrocyte known to remove myelin debris, apoptotic cell, and synapses^22^. Recently contribution of microglial phagocytotic ability to epileptogenesis has been reported^47,48^; therefore, phagocytotic activity of BCs may be involved with epileptogenesis in FCDIIb and mTORopathies. Given that we identified several other GO terms related to the possible function of BCs, in addition to the phagocytotic activity, further studies will be required to assess the physiological functions of BCs in FCDIIb and mTORopathies.

In conclusion, our study defines the molecular hallmarks of DNs and BCs in FCDIIb, providing mechanistic insight into their contribution to epileptogenesis. By generating the very first iST dataset of FCDIIb together with patient-matched snRNA-seq and proteomics data, we have provided a unique resource for advancing the study of mTORopathies. Our findings not only refine the mechanistic framework of epileptogenesis but also highlight potential therapeutic strategies targeting the molecular features of DNs and BCs in drug-resistant epilepsy.

## Methods

### Patient selection

In this study, patients (1) who had undergone neurosurgery for drug-resistant focal epilepsy at the National Center Hospital, National Center of Neurology and Psychiatry, Kodaira, Tokyo, Japan (NCNP), and (2) who were diagnosed with FCD, TSC or HME by the histopathologist were included. Temporal tip region of patient who had temporal lobe epilepsy with hippocampal sclerosis and underwent anterior temporal lobectomy were used as a control group. Clinical information and specimens were obtained according to the Declaration of Helsinki with written informed consent. This study was approved by the NCNP Ethics Committee, Japan (NCNP-A2018-050).

For Xenium experiments, the resected brain samples were fixed in 4% PFA for 48hrs, dehydrated in a graded alcohol series, cleared in xylene, and embedded in paraffin.

### Genome analysis

Multiple primer sets covering exonic and exon-intron border regions (+25 to −25) of genes involved in PI3K/AKT/mTOR signaling were designed using the Ion AmpliSeq Designer software (Thermo Fisher Scientific, Waltham, MA). The coverage rates of the targets were 90%. DNA derived from frozen tissues (20 ng) was amplified via polymerase chain reaction (PCR) using a premixed AmpliSeq HD Library Kit (Thermo Fisher Scientific) following the manufacturer’s protocol. Each library containing 50 pM was loaded on an Ion Chef™ Instrument (Ion Torrent™; Thermo Fisher Scientific). Prepared libraries were loaded onto Ion 540 Chips (four samples/chip) and sequenced with an Ion GeneStudio S5 System (Thermo Fisher Scientific), programmed for a read length of 200 bp and 500 flow cycles.

### Xenium spatial transcriptomics

Xenium in situ expression analysis was performed using Xenium Slides and Sample Prep Reagents (PN-1000460, 10X Genomics). FFPE samples were sectioned at 5 µm thickness, and placed on the Xenium slides. Deparaffinization and decrosslinking of sections were performed according to the manufacturer’s instructions (CG000580, 10X Genomics). Probe hybridization, ligation, and amplification were performed according to the user guide of “Xenium In Situ Gene Expression” (CG000749, 10X Genomics). Xenium Human Brain Gene Expression Panel (PN-1000599, 10X Genomics) was used. After washing, cell segmentation staining was performed using the Xenium Cell Segmentation Staining Reagents Kit (PN-1000639, 10X Genomics) overnight at 4°C. Autofluorescence quenching and nuclei staining were performed in the dark. The instrument was run using Xenium Analyzer (10X Genomics). To perform the post-Xenium staining, the slides were incubated in the quencher removal solution (10 mM sodium hydrosulfite) for 10 min, and washed in Milli-Q water and PBS according to Demonstrated Protocol (CG00613, 10X Genomics). The sections were stained with anti-nonphosphorylated neurofilament (SMI32, Biolegend, SMI32-R, 1:1,000) and anti-pS6 (CST, #5364, 1:800) antibodies. Following fluorescence imaging, the sections were re-stained with hematoxylin and eosin. Images were acquired using the NanoZoomer S60 digital whole slide scanner (Hamamatsu Photonics).

### Xenium data analysis

Cell segmentation was automatically performed with 10X nuclear and cell segmentation results in the Xenium Analyzer. The Xenium data were processed using Seurat pipeline (v5.3.0). The data were filtered based on the following parameters: nCount_Xenium > 10 and nFeature_Xenium >3. The filtered data were normalized using the NormalizeData in the Seurat package. Principal component analysis (PCA) was performed to reduce dimensionality using the top 20 PCs, and a shared nearest neighbor (SNN) graph was constructed using FindNeighbors. Cell clusters were identified using FindClusters (resolution = 0.5), and the results were visualized by uniform manifold approximation and projection (UMAP) with RunUMAP. Cell-type annotation was performed based on the expression of marker genes shown in Fig.1c. EN and AS clusters were sub-clustered using the top 30 PCs to reduce dimensions and a resolution of 0.5 to identify the clusters. EN sub-clusters were annotated using both marker genes and spatial information. Significantly expressed genes in sub-clusters were identified using FindMarkers function in the Seurat. Overlaying immunohistochemical and H&E images was performed on Xenium Explorer (v3.2.0) based on nuclear staining images.

### Nuclear isolation and snRNA-seq library preparation

Nuclear extraction protocol for human tissue was previously described^49^. Briefly, a frozen tissue was dounce homogenized in cold-lysis buffer (10 mM Tris-HCl pH8.0, 0.32 M Sucrose, 5 mM CaCl_2_, 3 mM Magnesium acetate, 2 mM EDTA, 0.5 mM EGTA, and 1 mM DTT, 0.1% Triton X-100). The homogenates were filtered with 100 µm and 40 µm strainers. To remove debris and leaking RNA, the homogenates were washed 3 times with wash buffer (10 mM Tris-HCl pH8.0, 0.32 M Sucrose, 5 mM CaCl_2_, 3 mM Magnesium acetate, 2 mM EDTA, and 0.5 mM EGTA) followed by centrifugation for 5 min at 500 x g at 4°C. The nuclear fraction was stored in storage buffer (10 mM Tris-HCl pH7.2, 0.43 M Sucrose, 70 mM KCl, 2 mM MgCl_2_, and 5 mM EGTA). snRNA-seq libraries were generated using 10X Chromium Next GEM Single Cell 3’ Reagent Kits v3.1 according to the manufacturer’s instructions. About 10000 nuclei per sample were loaded to generate libraries. The libraries were sequenced by NovaSeq6000 (Illumina).

### snRNA-seq data analysis

The CellRanger software (10X genomics) was applied to align the sequences to the pre-build human genome reference(refdata-gex-GRCh38-2020-A) containing GRCh38, and unique molecular identifiers (UMIs) were counted to construct the gene-barcode expression matrices. The matrices were processed using the Seurat pipeline. For quality control, potential doublets were detected using DoubletFinder (v2.0.6), and ambient RNA contamination was estimated using DecontX in Celda package (v1.22.1). To remove low-quality nuclei, the obtained data was filtered based on the following parameters: nFeature_RNA > 200, Doublet == “Singlet”, percent.mt < 5, and decontX_contamination < 0.2. The filtered data were normalized using the NormalizeData in the Seurat package. The normalized data were integrated using the IntegrateLayers (integrated.hm). PCA was performed to reduce dimensionality using the top 30 PCs, and a SNN graph was constructed using FindNeighbors. Cell clusters were identified using FindClusters (resolution = 1), and the results were visualized by UMAP with RunUMAP. Cell-type annotation was performed based on the expression of marker genes shown in Fig.5b. EN and AS clusters were sub-clustered using the top 30 PCs to reduce dimensions and a resolution of 1 to identify the clusters. Gene module score was calculated using AddModuleScore in the Seurat. Significantly expressed genes in each sub-cluster were identified using FindAllMarkers function in the Seurat. Functional enrichment of KEGG pathways and GO terms was performed using clusterProfiler (v4.14.6) and org.Hs.eg.db (v3.20.0). Reactome pathway analysis was performed on the Pathway browser (v3.7).

### LC/MS-MS analysis of human brain specimen

Frozen brain sections were powdered by using cyro-press CP-100WP (Microtech Nition) according to the manufacturer’s instructions. Powdered samples were lysed with 50mM Tris-HCl, 8M urea, 0.005% bromophenol blue, 1% SDS and sonicated for 20 sec. Protein concentration of each sample was measured by BCA assay (Fujifilm).

5 mg proteins were purified and digested by SP3 protocol (Hughes et al., 2019). Protein was reduced with 10 mM dithiothreitol at RT for 30 min and alkylated with 20 mM iodoacetamide at RT for 30 min in the dark. Samples were mixed with 150 μg of magnet beads (a 1:1 mixture of hydrophilic and hydrophobic SeraMag carboxylate-modified beads). Ethanol was mixed to a final concentration of 50%. The samples were mixed by a ThermoMixer C (Eppendorf, Hamburg, Germany) at 1200 rpm for 5 minutes at 24°C. The protein-bound beads were washed 3 times with 80% ethanol by using a magnetic rack.

Beads were resuspended with 100 mL of 50 mM ammonium bicarbonate. Trypsin/Lys-C Mix (1:50, w/w) was added to the sample tubes. Protein digestion was performed at 37°C with continuous mixing at 1,300 rpm overnight in a ThermoMixer C. Digested peptides were collected and desalted by using GL-Tip-SDB and resuspended in 20μL of 2% acetonitrile, 0.1% TFA solution.

Liquid chromatography-mass spectrometry(LC/MS) was performed on a Vanquish Neo UHPLC system (Thermo Scientific) coupled with Orbitrap Fusion (Thermo Scientific). Peptide samples were loaded onto a trap column (Acclaim PepMap 100 C18 LC column, 3 μm, 75 μm ID × 20 mm; Thermo Fisher Scientific) and were separated on an EASY-Spray C18 LC column (75 µm × 15 cm, 3 µm, 100 Å, Thermo Fisher Scientific) by a linear gradient consisting of 0%-35% acetonitrile with 0.1% formic acid over 120 min at a flow rate of 300 nL/min.

In data-independent acquisition (DIA) method, MS1 scan range was set at 390-1010Lm/z in positive mode, orbitrap resolution at 120,000, and standard AGC target mode with maximal injection time of 60 ms. MS2 scan range was set 400-1001 m/z with orbitrap resolution at 60,000 and normalized HCD collision energy was set at 30%. Number of scan event was 50 with 12 m/z isolation windows in quadrupole mode.

DIA data was processed using DIA-NN (v1.9.2). Default settings for in-silico spectral library preparation were used. A human protein database from UniprotKB (UP000005640_2024_10_18.fasta) was used to prepare an in-silico spectral library. DIA-NN analysis was performed using default settings (Trypsin/P digestion, one missed cleavage is allowed, carbamidomethylation on cysteine as fixed modification, acetylation on the N-term as variable modification, false discovery rate for peptides and proteins were set to 0.01, match between runs was enabled).

### Immunohistochemistry

FFPE samples were sectioned to 6 µm thickness. Immunohistochemical staining was performed as previously described ^50^. Briefly, the sections were incubated with 0.3% H_2_O_2_ in methanol for 30Lmin to inactivate the intrinsic peroxidase. Antigen retrieval was carried out by autoclaving for 10Lminutes in 10LmM sodium citrate buffer at 121°C.

The specimens were incubated overnight at 4°C with primary antibodies diluted with 5% normal goat serum in PBS. Immunoreaction products were detected by the polymer immunocomplex method using the Envision system (Dako) and visualized using 3, 3′-diaminobenzidine (DAB). Sections were then counterstained with hematoxylin. The following primary antibodies were used: anti-CROT (Proteintech, 1:200) and anti-CAV1 (Proteintech, 1:500).

## Data availability

Raw sequence files and spatial transcriptomic data obtained in this study will be available after publication. Before that, all data will be shared upon reasonable request.

## Code availability

Processing workflows used for all analysis are available in Github (https://github.com/Hoshino-lab/FCD_Shimaoka_2025)

## Acknowledgments

We are greatly appreciating the patients and their families for participating in this study.

Illustration in the figures were created with BioRender.

## Contributions

K.S., S.M., K.I., M.I., and M.H. conceived and designed the study. K.I. and M.I. coordinated patient recruitment and neurosurgical tissue collection. K.S., S.M., N.N.K.T., K.H., N.A., S.H., M.M., K.N., and T.O. contributed to transcriptome data generation and omics data analysis. K.Y., E.U., T.S., and M.T. contributed to tissue processing and immunohistochemistry experiments. S.T., T.N., and K. Kaibuchi contributed to proteomics data generation. K.I., K.M., and K. Komatsu performed genome analysis. K.S. and S.M. wrote the original draft. M.S., M.I., and M.H. reviewed and edited the manuscript. All authors reviewed and approved the final manuscript.

---

**Figure S1.**
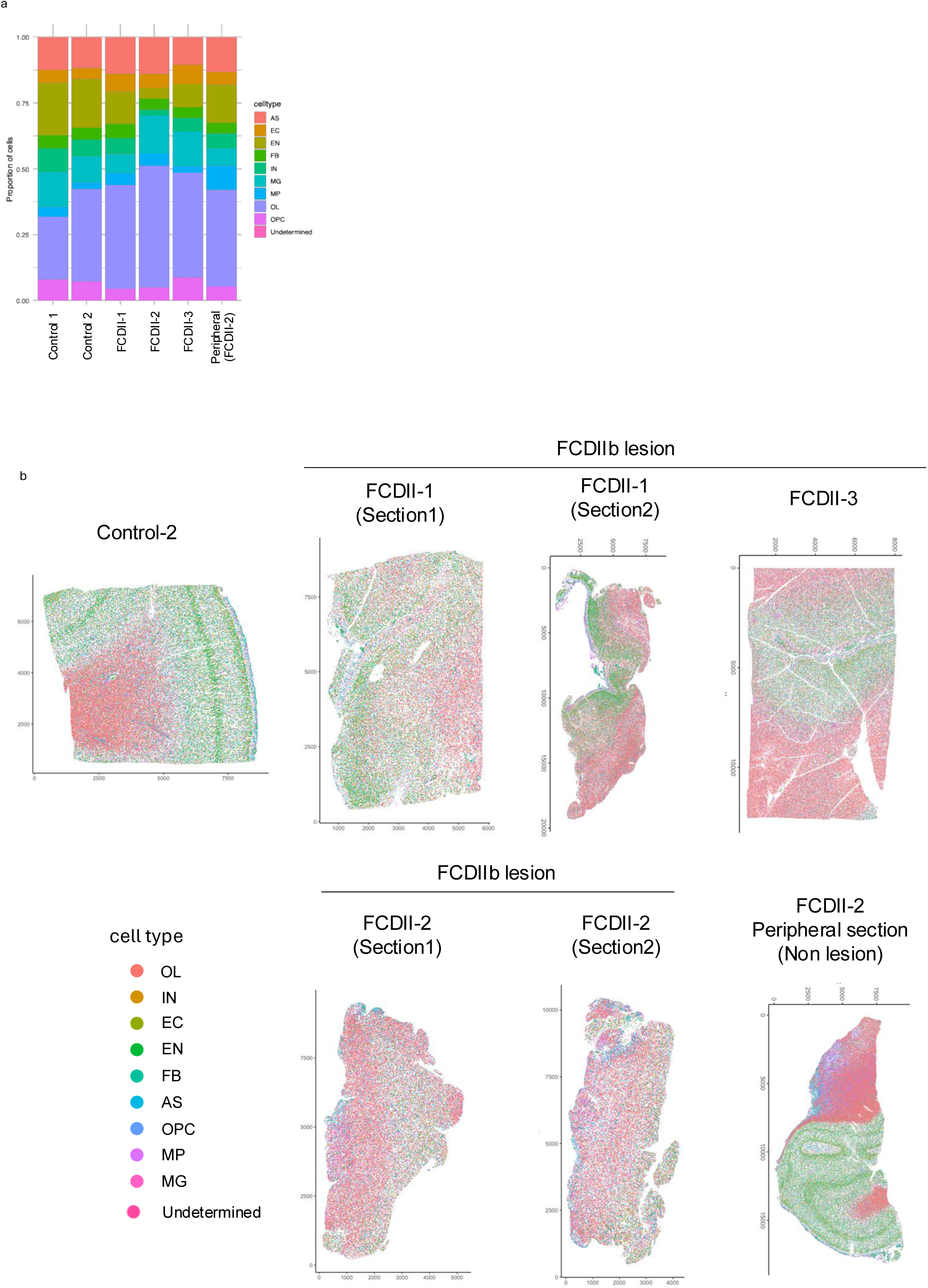
Spatial distribution of major cell classes across samples. (a) Proportional composition of major cell type clusters across samples. (b) Spatial maps of annotated cell types across controls and FCDIIb sections (lesion and peripheral non-lesion tissue).

**Figure S2.**
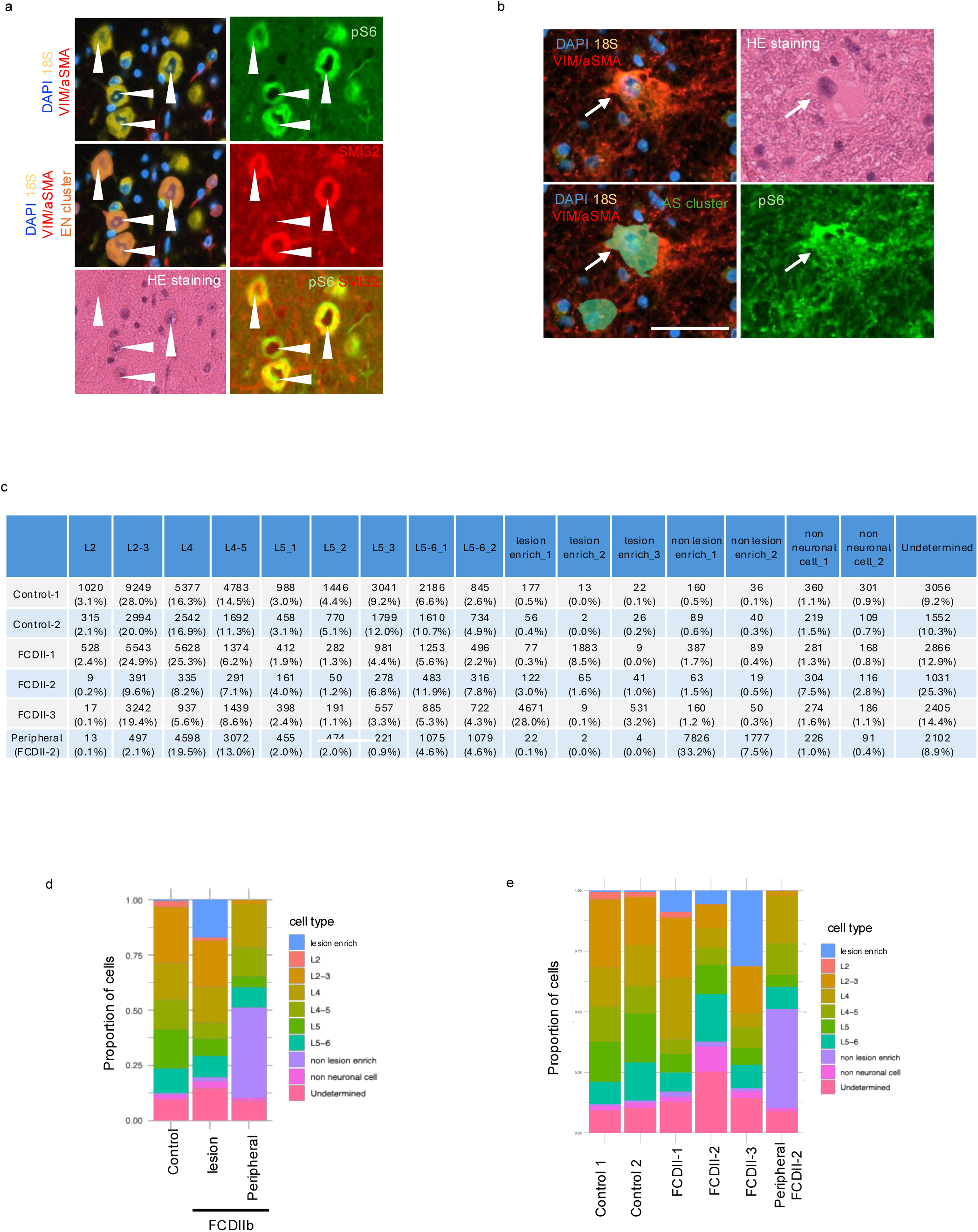
Quantification of neuronal cluster composition and immunofluorescence examples. (a,b) Cluster identity validation using marker staining (pS6 and SMI32) together with boundary staining (DAPI, 18S, VIM/αSMA). pS6/SMI32 double-positive DNs mapped to excitatory neuron (EN) clusters (a), whereas pS6-positive BCs mapped to astrocyte (AS) clusters (b). Scale bars; 50 µm. (c) Table summarizing excitatory neuron subcluster counts and percentages for each (d,e) Proportional composition of major cell type clusters across groups (d) and samples (e). sample.

**Figure S3.**
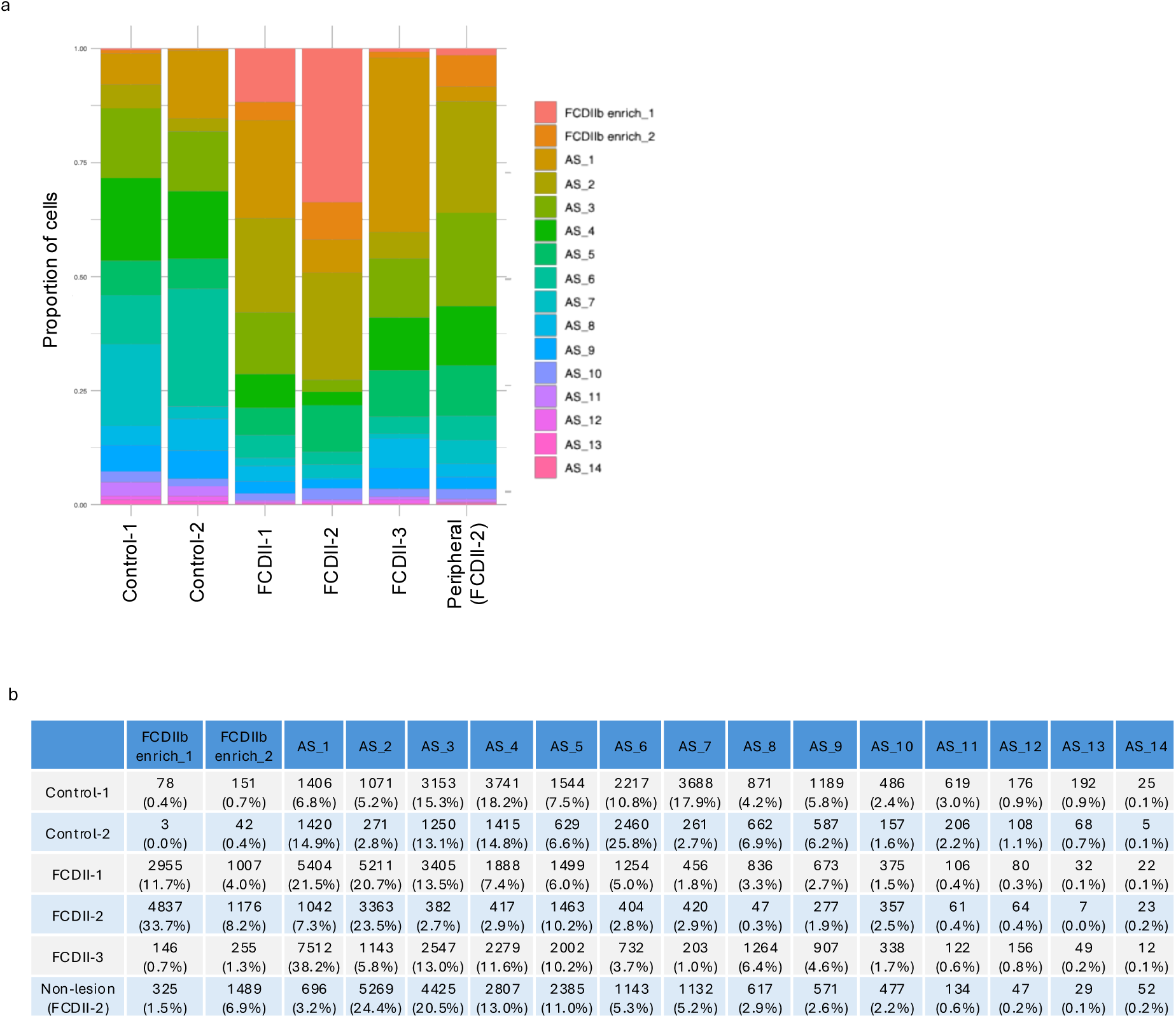
Astrocyte subcluster composition across samples. (a) Proportional composition of astrocyte subclusters across samples. (b) Table summarizing astrocyte subcluster counts and percentages for each sample.

**Figure S4.**
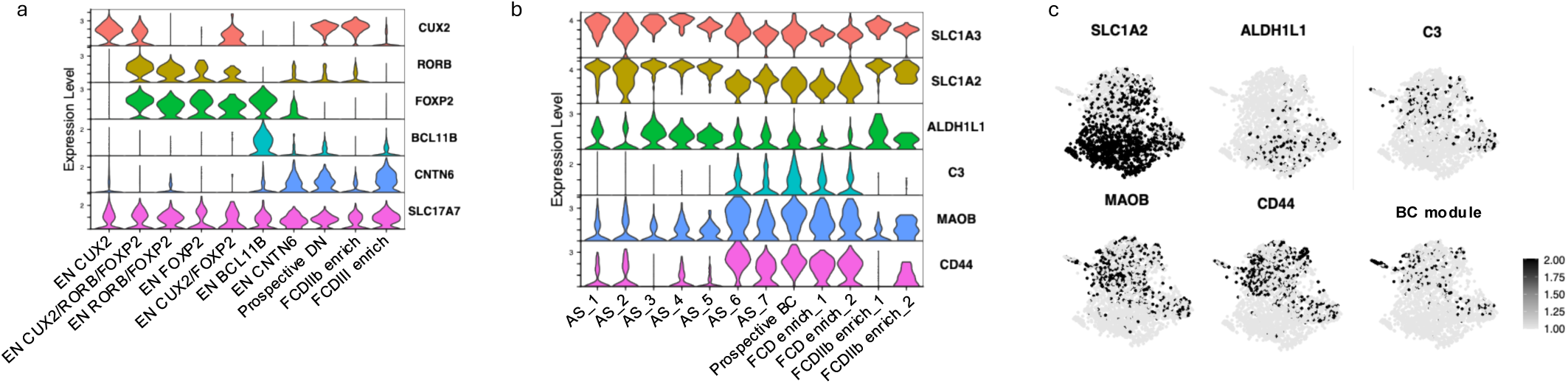
Marker expression and DN and BC module scores in EN and AS subclusters. (a) Violoin plot showing expression of pan-excitatory neuron marker *SLC17A7* and layer markers (*CUX2, RORB, FOXP2, BCL11B, CNTN6*) across predicted prospective DN cluster and EN subclusters. (b) Violoin plot showing expression of pan-astrocyte markers (*SLC1A3, SLC1A2, ALDH1L1*) and reactive astrocyte markers (*C3, MAOB, CD44*) across predicted prospective BC cluster and AS subclusters. (c) Feature plots showing spatial distribution of selected astrocyte markers and BC module score.

**Figure S5.**
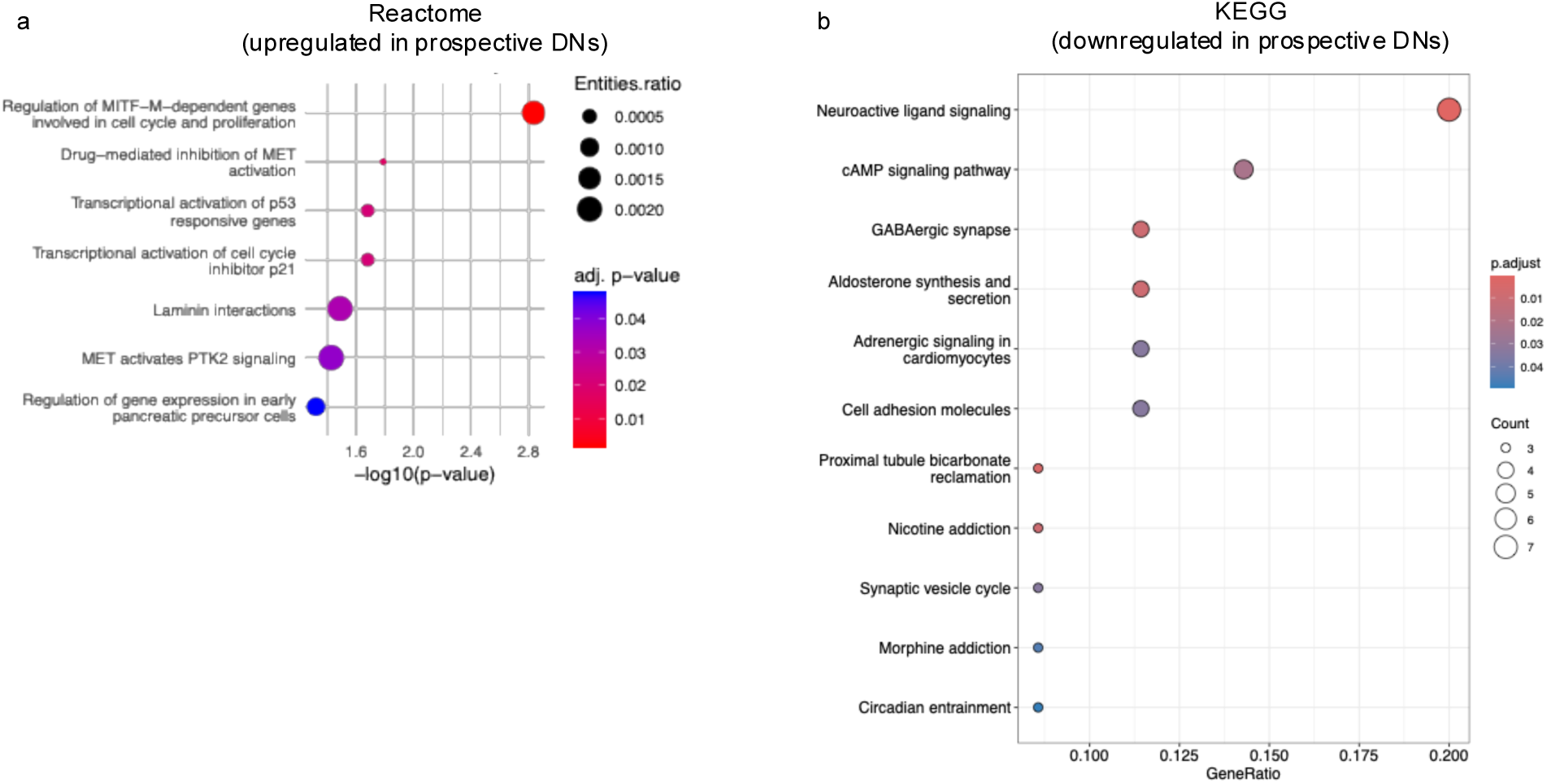
Reactome and KEGG enrichment analyses for prospective DNs. (a) Reactome pathway enrichment of genes upregulated in prospective DNs. (b) KEGG pathway enrichment of genes downregulated in prospective DNs.

**Figure S6.**
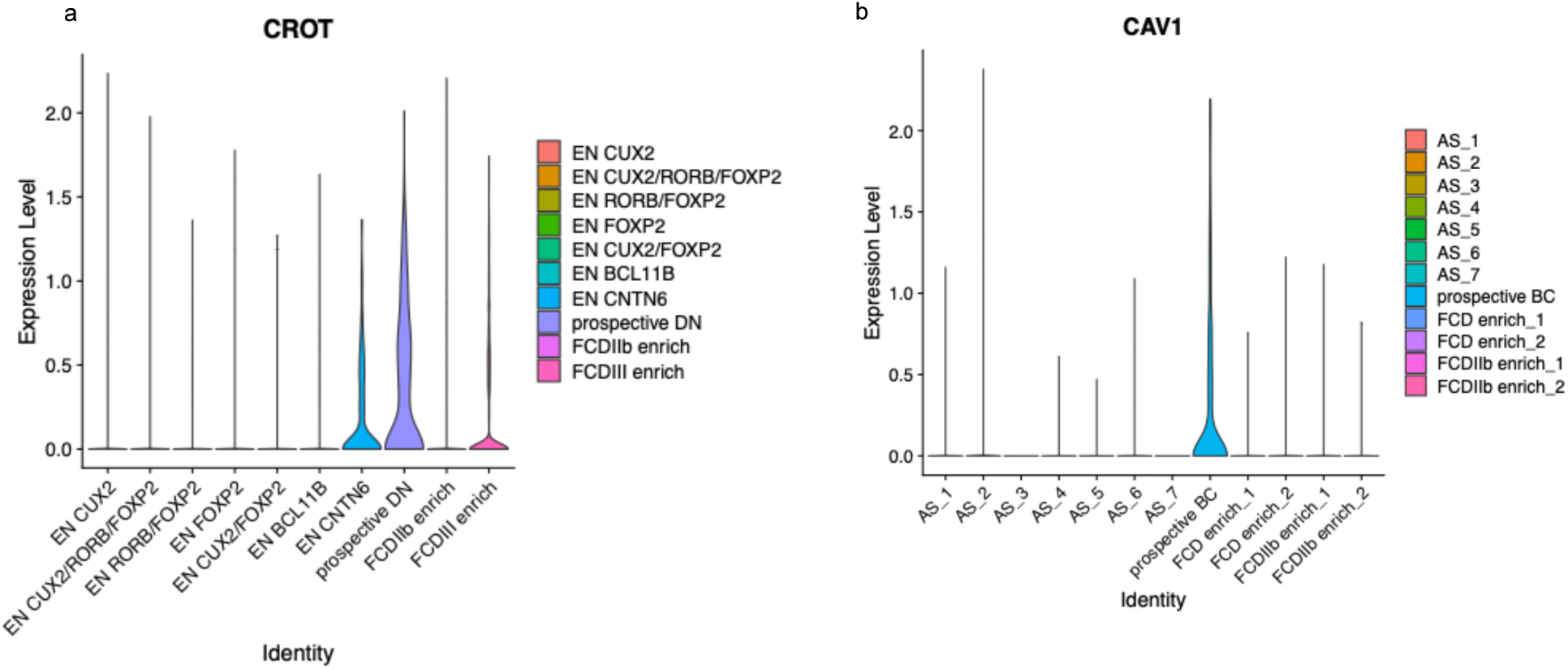
Expression of candidate marker genes for DNs and BCs. (a,b) Violin plots showing expression levels of CROT across and prospective DN cluster and EN subclusters (a), and CAV1 across prospective BC cluster and AS subclusters.

